# Complementary Maps for Location and Environmental Structure in CA1 and Subiculum

**DOI:** 10.1101/2021.02.01.428537

**Authors:** Jacob M Olson, Alexander B Johnson, Lillian Chang, Emily L Tao, Xuefei Wang, Douglas A Nitz

## Abstract

The dorsal subiculum lies among a network of interconnected brain regions that collectively map multiple spatial and orientational relationships between an organism and the boundaries and pathways composing its environment. A unique role of the subiculum in spatial information processing has yet to be defined despite reports of small neuron subpopulations that encode relationships to specific boundaries, axes of travel, or locations. We examined the activity patterns among populations of subiculum neurons during performance of a spatial working memory task performed within a complex network of interconnected pathways. Compared to neurons in hippocampal sub-region CA1, a major source of its afferents, subiculum neurons were far more likely to exhibit multiple firing fields at locations that were analogous with respect to path structure and function. Subiculum neuron populations were also found to exhibit a greater dynamic range in scale of spatial representation and for persistent patterns of spiking activity to be aligned to transitions between maze segments. Together, the findings indicate that the subiculum plays a unique role in spatial mapping, one that complements the location-specific firing of CA1 neurons with the encoding of emergent and recurring structural features of a complex path network.

## Introduction

The dorsal subiculum is situated within a distributed system of brain regions forming a ‘cognitive map’ that encodes an organism’s spatial relationship to its environment (Andersen et al., 1973; Hjorth-Simonsen, 1973; Amaral & Witter, 1989; Amaral et al., 1991; Witter, 2006). The few studies that have examined the impact of damage selective to subiculum (SUB) support a role for this structure in the two most prominent tests for spatial navigation, the Morris water tank and T-maze spatial alternation tasks (Morris et al., 1990; Galani et al., 1998; Frost et al., 2020). Despite this, relatively few studies have addressed the form or forms by which subiculum neurons represent spatial and orientational relationships to the environment.

Clues to a unique role for SUB as a component of a distributed cognitive mapping system come from its major sources of afferents (Witter, 1990; Witter et al., 2000; Naber et al., 2001; Cembrowski 2018). A prominent input from the anterior thalamus and moderate input from the presubiculum suggests a strong influence of orientation or ‘head direction’ tuning on SUB function (Goodridge & Taube, 1997; Winter et al., 2015; Viejo & Peyrache, 2019; O’Mara & Aggleton, 2019). Indeed, sensitivity to head orientation relative to environmental boundaries was observed in early studies examining the firing of SUB neurons during random foraging in open arenas (Barnes et al., 1990; Sharp & Green, 1994; Sharp, 1999) and, more recently, in a track based environment (Olson et al., 2017). Major inputs from both medial and lateral entorhinal cortex and hippocampal sub-region CA1 indicate a strong influence of tuning by location within an allocentric, ‘world-centered’, space defined by environmental boundaries (Muller & Kubie, 1987; Hafting et al., 2005). Accordingly, evidence for ‘place-specific’ firing (Sharp & Green, 1994; O’Mara, 2005; Kim et al., 2012; Olson et al., 2017; Lee et al., 2018) in SUB has been observed in a small percentage of neurons as has encoding of position and orientation relative to boundaries and barriers (Lever, 2009; Stewart et al., 2014; Stensola et al., 2015; Olson et al., 2017; Viejo & Peyrache, 2019; Poulter et al., 2020). Finally, SUB appears to play a role in the encoding of objects and landmarks and their relationships to environmental boundaries (Sun et al., 2019; Poulter et al., 2020).

Several studies examining the spatial firing properties of SUB neurons compared them to neurons of the CA1 sub-region of hippocampus, a region well-known for the presence of ‘place cells’ (O’Keefe & Dostrovsky, 1971). From such work, it is known that SUB neurons are less focused in their tuning to environmental location, tend to exhibit a greater number of individual firing fields, and have a greater tendency to show similar spatial firing patterns across two arena environments that are the same but for differences in shape (circle versus square arena) or size (Sharp & Green, 1994; Sharp, 1997; Sharp, 1999 (2); Kim et al., 2012; Olson et al., 2017). These quantitative differences in tuning may function to permit generalization across similar environments or to maximize the efficiency of information output to efferent targets (Sharp, 1999; Kim et al., 2012). Nevertheless, it remains to be determined whether spatial tuning in SUB and CA1 can be viewed as part of a continuum or whether substantive differences in tuning evidence different contributions of each region to components of a distributed cognitive map.

To draw out potential differences between SUB and CA1, we examined the spatial firing properties of neuron populations in these regions during performance of a complex spatial working memory navigational task set within a complex network of pathways. The combined structure of the path network and set of task rules allowed us to determine to what extent rats exhibited behavior consistent with having knowledge of the overall layout of individual path segments and their relation to the full path network. The routes utilized to meet task demand bore structural similarity in having the same total number of turns and distances between them. This allowed us to detect spatial tuning of neurons that reflect emergent properties of the combined task and maze structure. Partial overlap between routes allowed us to determine whether trajectory-dependence (Frank et al., 2000; Wood et al., 2000; Ferbinteanu & Shapiro, 2003; Ainge et al., 2007) differs between CA1 and SUB and whether its presence or absence is a dynamic property of either system. Finally, the maze structure contained many intersections and path segments of different lengths, allowing us to compare the spatial scale of representation for CA1 versus SUB as well as the alignment of their population firing patterns to maze structure.

During sustained, highly accurate performance of the task, we find that individual neurons of the CA1 region primarily encode the animal’s location and the trajectory taken through that location. In contrast, just one synapse beyond CA1, SUB neurons were often active for “kinds” of places that were analogous with respect to maze structure and with respect to multiple spatial variables such as head direction, axis of travel, and progression through a route. We also find that the spatial scale of representation is greater for SUB, and that it varies dynamically in both structures. Furthermore, persistence in CA1 and SUB population activity patterns across track positions segmented space in distinct ways, suggesting that spatial representation in these structures can follow qualitatively different rules. These findings identify a unique role for SUB in spatial cognition and suggest that this region is critical to encoding the fundamental structure of pathways through complex environments.

## Results

### Robust navigation within a complex environment

We trained 6 rats on a variation of the “Triple-T” maze (Figure 1A, Olson et al., 2017; Olson et al., 2020). The task demands that animals learn the functional relationships among a set of interconnected pathways and provides a means by which to assess the impact of location, orientation and trajectory on the spiking activity of recorded neurons. Briefly, on each trial, the animal must move through 3 sequential left or right turns to arrive at 1 of 4 goal locations. The rat must then return to the starting location via external pathways that surround the internal pathways. Rewards at each goal location were 1/4 Cheerio piece and distributed on a “visit-all” schedule where reward access reset after all locations were visited. During post-implantation recording sessions, rats ran on average 148 trials per session.

**Figure 1:**
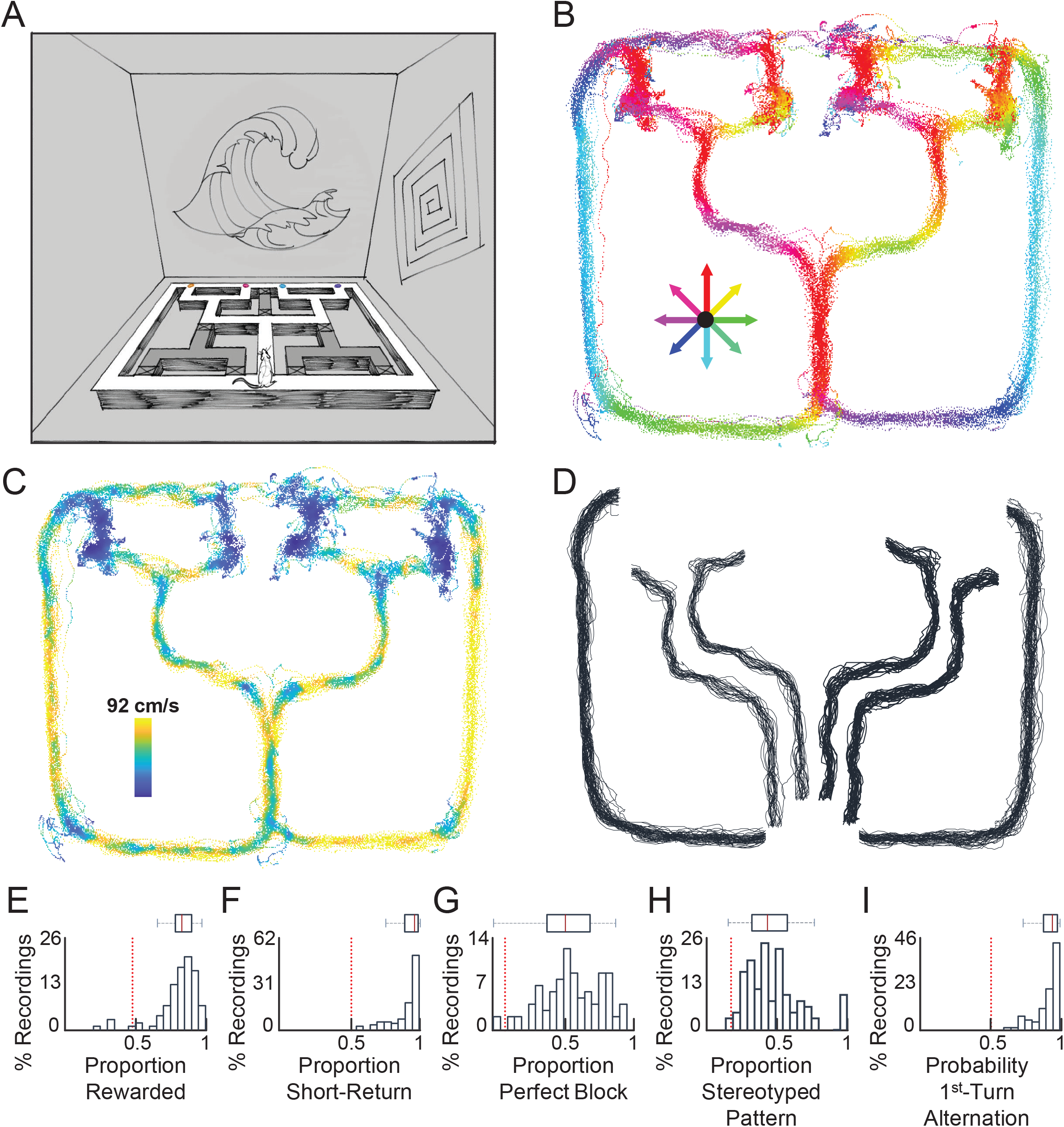
Robust Navigation of a Complex Environment. **A)** Illustration of Triple-T navigational task recording environment. Animals begin at a starting point (center-bottom) and must navigate a three-turn sequence to arrive at one of four different reward locations. In the visit-all-4 reward schedule, rats must visit each of the four locations before revisiting any reward location. **B)** Behavioral tracking from a full sample recording on the Triple-T maze. Positional data is color-coded showing the animal’s average head direction at each position. **C)** Behavioral tracking, as in B, but with color coding showing average velocity, from 0 in blue to the max, 92cm/s, in yellow. **D)** Behavioral tracking of identified routes. Shown are all data labeled as uninterrupted runs from the same recording session in the visit-all-4 reward schedule. In this setup, four outbound routes begin at the start point and end at four different reward locations. Two external routes are also defined encapsulating the paths from the rewards to the starting location. Routes are minimally translated and stretched for visualization purposes. **E-I)** Box plots of per behavioral session performance metrics *(red bar: median; box limits: first and third quartiles; whiskers: range of non-outlier data points)*. In each, the dotted red line represents chance. **E)** Proportion of rewarded runs. **F)** Proportion of shorter return routes taken. **G)** Proportion of visit-all-4 reward blocks without a mistake. **H)** Proportion of the most commonly run block pattern from each recording. **I)** Proportion of left/right alternations at the first choice point.

The nature of this task and our training regimen purposely leads to highly stereotyped running behavior. The rats became proficient in the procedural aspect of the task, only moving in one direction at any given location (Figure 1B) and doing so at high speeds (Fig 1C). For analysis purposes, we selected out behavioral epochs where the animal ran uninterrupted from the start point to one of four reward locations (outbound routes) or from the reward locations back to the starting point (return routes, Fig 1D). This results in analyzing only the behavior during locomotion and controls over the range of head orientations and actions associated with any specific location. The rats met the uninterrupted running criteria on 85% of route traversals and averaged a velocity of 57 cm/sec during the accepted runs.

Rats also became proficient at the working memory aspect of the visit-all task on the Triple-T maze, receiving rewards on 81% of trials in recordings post-surgery (Figure 1E, mean 0.81 ± 0.15 s.d.; N = 95, P < 0.0001, one-sided Mann-Whitney U test; chance = 48%). The animals exhibited an understanding of the track space, taking the shorter return pathway on 92% of trials (Figure 1F, mean 0.92 ± 0.11 s.d.; N = 95, P < 0.0001, one-sided Mann-Whitney U test; chance = 50%). Animals even ran perfect blocks – a series of visiting all 4 locations without mistake – on 56% of trials (Figure 1G, mean 0.56 ± 0.21 s.d.; N = 95, P < 0.0001, one-sided Mann-Whitney U test; chance = 9%). Performance at this level may suggest formation of a habit with respect to the order of internal routes taken over a block, but a closer look at animal tendencies disagrees. The most common pattern used by an individual animal over an individual recording session accounted for only 48% of correct blocks (Fig 1H, mean 0.48 ± 0.20 s.d.; N = 94, P < 0.0001, one-sided Mann-Whitney U test; chance = 19%), and 26% ± 14% of total blocks. However, animals clearly used alternation at the first turn location as a mnemonic strategy, doing so on 91% of trials (Fig 1I, mean 0.91 ± 0.09 s.d.; N = 95, P < 0.0001, one-sided Mann-Whitney U test; chance = 50%).

### Individual subiculum neurons are active across analogous maze locations

To compare neural activity in SUB and CA1, we recorded from the same 6 animals during task performance using microdrives outfitted with tetrodes. 573 neurons were recorded from dorsal SUB (4 SUB animals), with the majority being from the proximal half of SUB, and 401 neurons were recorded from dorsal CA1 (3 CA1 animals) (Figure 2B, Supplemental Figure 1). Maximum and minimum firing rate thresholds were used to exclude inactive cells and putative interneurons, leaving a final dataset of 480 SUB and 298 CA1 neurons.

**Figure 2:**
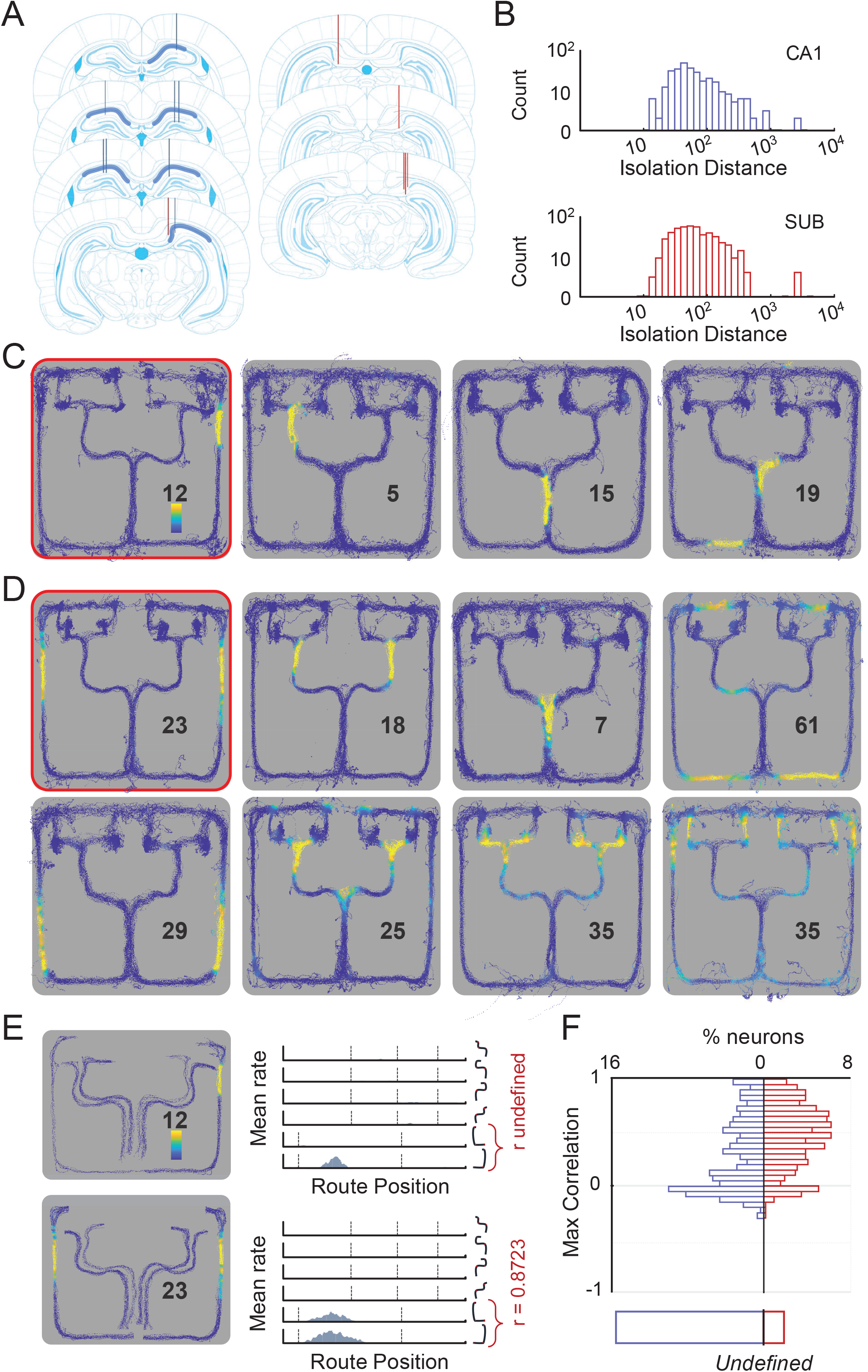
Individual Subiculum Neurons Are Active in Analogous Spaces. **A)** Electrode placement in CA1 (blue, 3 rats) and SUB (red, 4 rats). Lines end at the terminal identified locations of tetrode bundle tracks. **B)** Isolation distance waveform discrimination quality metric for both CA1 (top) and SUB (bottom). **C)** Positional firing rate maps of example CA1 neurons, color mapped from 0 (blue) to the mean + 3 s.d. (yellow) for each neuron. The max value is written in each map. **D)** Positional firing rate maps of example SUB neurons, colormapped as in **C. E)** *Left:* Two dimensional spatial maps of mean firing rates for individual neurons highlighted in **C** and **D**, mapped as a function of route and track position. Like **Figure 1C**, routes are minimally translated and stretched from the actual track location to separate each map for visualization purposes. Colormap scaling is identical to **C**. *Right:* Linearized mean firing rates as a function of routes for the corresponding neurons on the left. The routes for each graph are depicted on the right. Pearson correlations of the two return routes are shown. When one route has no activity, correlations are undefined (top). **G)** Population histogram of maximum pairwise correlations of routes for individual cells. If only one route had activity, the correlation was undefined.

Commonly used measures for spatially tuned CA1 and SUB spike firing were largely consistent with previous literature. SUB neurons fire more (CA1 mean 0.89 ± 1.43 s.d. N = 298; SUB mean 3.62 ± 4.23 s.d. N = 480, P < 0.0001, one-sided Mann-Whitney U test) and exhibit lower spatial information per spike (CA1 mean 2.97 ± 1.18 s.d. N = 298; SUB mean 1.55 ± 1.21 s.d. N = 480, P < 0.0001, one-sided Mann-Whitney U test) (Kim et al., 2012) as well as lower spatial selectivity (CA1 mean 63.9 ± 50.8 s.d. N = 298; SUB mean 31.1 ± 38.8 s.d. N = 480, P < 0.0001, one-sided Mann-Whitney U test) (Kim et al., 2012, Lee et al., 2018). There was no difference between CA1 and SUB in spatial coherence on the maze (CA1 mean 0.49 ± 0.12 s.d. N = 298; SUB mean 0.49 ± 0.14 s.d. N = 480, P = 0.82, Mann-Whitney U test), unlike previous reports (Sharp & Green, 1993, Lee et al., 2018).

The differences in general spatial properties were qualitatively apparent in CA1 and SUB activity on the Triple-T maze. Individual SUB neurons often exhibited larger or more firing fields than their CA1 counterparts. Beyond these previously-described differences in spatial tuning, we observed that many SUB neurons displayed another differentiating trait: increased structural relationships between the firing fields of individual neurons. While some SUB neurons did show single fields on the track, activity of individual SUB neurons more often occurred in multiple locations that often shared spatial or functional features (Figure 2D). For simplicity, we will refer to this as representation of structural or functional analogy between maze locations.

To quantify this propensity for multiple firing fields of individual neurons to distribute across analogous maze locations, we created linearized positional firing rate maps for each individual route. We then correlated individual neuron’s firing activity for non-overlapping portions of all combinations of outbound runs as well as between the two return runs. For neurons with fields at analogous locations along two or more routes, correlations should be high, whereas correlations will be low or negative if the locations of firing fields across analogous portions of two routes are very different (Figure 2E). Rate vector correlations are mathematically undefined in cases where a neuron does not fire at all along one of two routes, but we note that, practically speaking, this can be considered a low or zero correlation result. We found that SUB neurons were much more likely to exhibit high correlations between routes. SUB neurons had higher maximum correlations between routes (CA1 mean 0.29 ± 0.34 s.d. N = 252; SUB mean 0.46 ± 0.30 s.d. N = 468, one sided KS test, P < 0.0001, KS test stat 0.29), as 47% of SUB neurons had either internal or external routes with correlations exceeding 0.5, as compared to 26% of CA1 neurons (Figure 2F). Thus, our simple measure to detect similarity in positional firing rates for analogous but spatially segregated locations along same-length routes provides a strong indication of a qualitatively different organization of spatial tuning for SUB versus CA1 neurons.

### Analogous responses in subiculum include decreased trajectory dependence

Since individual SUB neurons exhibit much greater similarity in firing rates across analogous spaces of two routes than CA1 neurons, we next determined to what extent this is also seen in the two regions’ population activity patterns. We first present data for the long straight segments of the return routes that have functional and directional similarity but are separated by nearly 2 meters (Figure 3A, labeled B). Pearson correlations of population mean firing vectors between the two segments show higher correlations for SUB than CA1 (Figure 3B, CA1 mean 0.28 ± 0.09 s.d. N = 110; SUB mean 0.60 ± 0.08 s.d. N = 110, P < 0.0001, one-sided Mann-Whitney U test). We observed low correlation in population activity patterns as compared to the correlations for odd versus even trials of the same location (All control comparisons, N = 110, P <0.0001, one-sided Mann-Whitney U test). The final straight segments on the return routes maintain functional similarity but lack directional equivalence (Figure 3A, labeled C). Yet, SUB again had higher correlations than CA1 (Figure 3C, CA1 mean 0.25 ± 0.07 s.d. N = 59; SUB mean 0.64 ± 0.07 s.d. N = 59, P < 0.0001, one-sided Mann-Whitney U test; all control comparisons, P <0.0001, one-sided Mann-Whitney U test), and the segments’ correlations for SUB were not measurably lower than the directionally consistent segments (P > 0.9984, one-sided Mann-Whitney U test).

**Figure 3:**
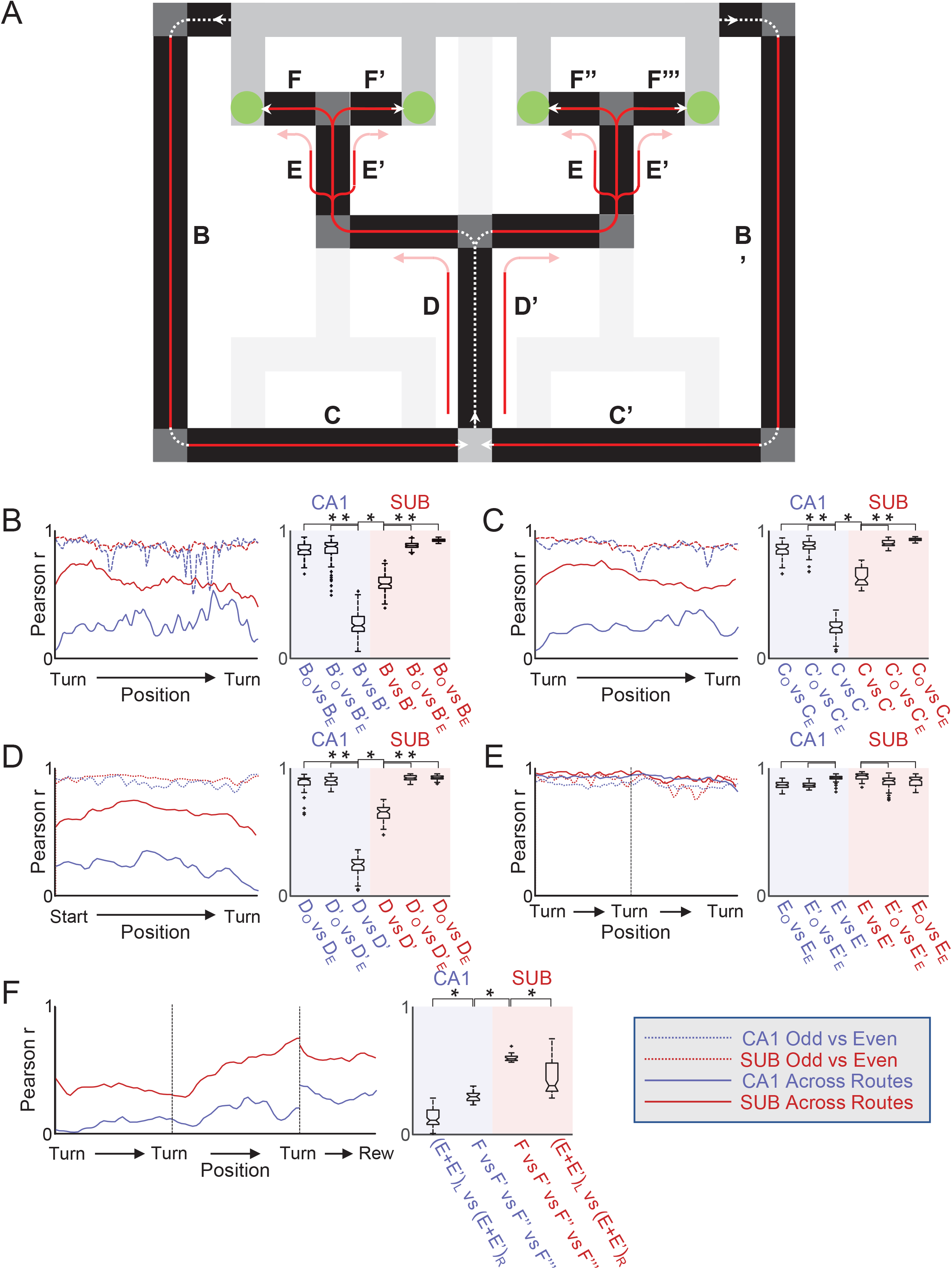
Analogous Responses in Subiculum Include Decreased Trajectory Dependence. **A)** Schematic of Triple-T maze. Highlighted regions are analyzed in the following panels corresponding with the labels. White dotted lines show animal routes, with solid red portions indicating regions analyzed and shown in this figure. **B-F)** *Left:* Pearson correlations of population mean firing vectors, and *Right:* box plots of the Pearson correlation distributions. In all panels, CA1 data is blue and SUB is red. Solid lines are across route segment comparisons. Dotted lines are odd versus even correlations within one group. For all box plots, black bar: median; notch: comparison interval at 5% level; box limits: first and third quartiles; whiskers: range of non-outlier data points; +: outliers). Subscript O is for odd trial population data, while E for even trial population data. * denotes Mann Whitney U test with P < 0.05. **B)** CA1 and SUB population correlations between the long straight segments of the two return routes. **C)** CA1 and SUB population correlations between the final straight segments of the two return routes. **D)** CA1 and SUB population correlations of the center stem split according to upcoming turn direction. **E)** CA1 and SUB population correlations of the two straight segments preceding the final outbound turn, split by upcoming turn direction. **F)** CA1 and SUB population correlations of non-overlapping outbound run segments.

We also used the population rate vector correlation technique to contrast pattern recurrence for SUB versus CA1 populations over overlapping portions of outbound routes. One well-described example of differentiation in CA1 is trajectory-dependent coding, wherein CA1 in-field firing rates vary according to the trajectories taken through the field (Frank et al., 2000; Wood et al., 2000; Ferbinteanu & Shapiro 2003, Ainge et al. 2007; Grieves et al., 2016). In the Triple T maze, the center stem (Figure 3A, labeled D) is a segment common to all four of the outbound routes. Correlation of mean firing rates between pre-left-turn and pre-right-turn activity show strong trajectory dependence in the CA1 population. CA1 population activity correlations were low as compared to the correlations for odd versus even trials of the same trajectory (Figure 3D, CA1 mean 0.24 ± 0.07 s.d. N = 51; both P < 0.0001, one-sided Mann-Whitney U test). The SUB population exhibits far less trajectory-dependence than the CA1 population (SUB mean 0.66 ± 0.06 s.d. N = 51; P < 0.0001, one-sided Mann-Whitney U test; all control comparisons, P < 0.0001, one-sided Mann-Whitney U test). That is, the SUB population activity patterns largely generalize in the representation of this space despite the very different trajectories. We note that trajectory dependence is nonexistent in both CA1 and SUB for the space where the animals approached the final pre-reward choice point (Figure 3A, labeled E, Figure 3E, CA1 mean 0.92 ± 0.02 s.d. N = 67; SUB mean 0.94 ± 0.03 s.d. N = 67, all control comparisons, P > 0.9999, one-sided Mann-Whitney U test). This indicates that trajectory-dependence is dynamic in its expression for CA1 and for SUB, but much lower overall for SUB.

Finally, we examined encoding similarity as the animal approaches the goal locations (Figure 3A, labeled F). We hypothesized that SUB may generalize across the approaches to these four separate locations. Transitioning to the final segment, spatial locations diverge. However, correlations across the routes increased for both CA1 and SUB (Figure 3F, CA1 pre-final segment mean 0.13 ± 0.07 s.d. N = 67, CA1 final segment mean 0.30 ± 0.04 s.d. N = 21, P > 0.0001, one-sided Mann-Whitney U test; SUB pre-final segment mean 0.45 ± 0.14 s.d. N = 67, SUB final segment mean 0.61 ± 0.03 s.d. N = 21, P > 0.0001, one-sided Mann-Whitney U test), with SUB again generalizing more (N = 21, P > 0.0001, one-sided Mann-Whitney U test).

### Subiculum and CA1 population firing patterns chunk epochs differentially relative to task phase and maze structure

Given the striking differences between SUB and CA1 encoding of multiple task phases, we looked deeper into how population firing patterns shift relative to task phase and maze structure. For each region, we assembled ensemble firing rate vectors of the even and the odd trials separately, using all recorded neurons at each position along each route. We then concatenated these route-based positional rate vectors and calculated Pearson correlations to assess pattern similarity for each route location relative to all others (Figure 4AB, Cowen & Nitz, 2012). The resulting correlation matrix can be used to compare how SUB and CA1 population activity patterns persist over contiguous locations and whether shifts in patterning are related to specific task phases or the beginnings and endings of maze segments.

**Figure 4:**
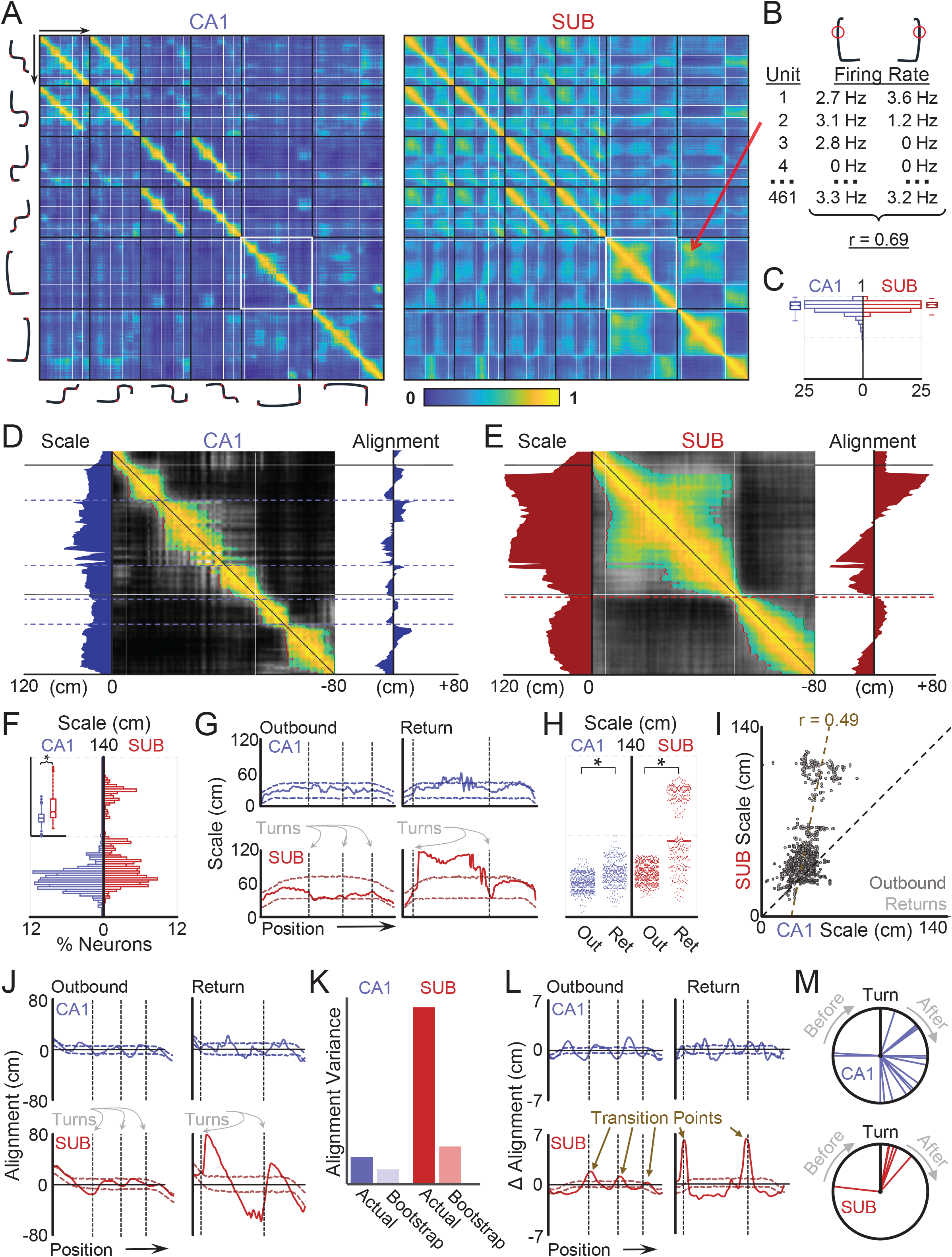
Subiculum and CA1 Populations Chunk Epochs at Different Locations. **A)** Odd versus even trials population correlation matrices for CA1 (left) and SUB (right). The vector of mean firing rates for all recorded cells in the given region are correlated across each pairwise position and route combination. High correlations (yellow) indicate similar encoding of the two spaces. The routes that are linearized are depicted next to the corresponding rows/columns. Dotted white lines indicate turn apexes. **B)** Example population correlation calculation. The matrix value indicated by the red arrow on **A** is the correlation of the odd or even mean firing rates from all neurons at the corresponding route and position locations. Here, it is two corresponding locations from the two return routes. The color mapped value is the Pearson correlation of these two ensembles. **C)** Histograms of odd versus even trials population correlations for the same locations. This corresponds to the diagonal values in the population correlation matrices. **D-E)** Representational scale and alignment examples. **D)** Center: Subsection of population correlation matrix highlighted in **A**. Contiguous row-wise off-diagonal values above 0.5 are mapped to color on the same scale from A. The remainder of the submatrix is mapped black to white. The width of the above threshold region (representational scale) is shown on the left along the route space. The difference in forward/backward extent of the above threshold region (representational alignment) is shown on the right. Scales are in cm. Black lines show turn apexes, while dotted blue lines show segmentation transition points. **E)** Same as D, but for SUB. Chunking edge lines are in red not blue. **F)** Histogram of representational scale of CA1 (left, blue) and SUB (right, red). Inset: box plots of same data (bar: median; notch: comparison interval at 5% level; box limits: first and third quartiles; whiskers: range of non-outlier data points; +: outliers). * denotes Kolmogorov-Smirnov test with P < 0.05. **G)** Mean representational scale for the outbound (left) and return (right) runs for both CA1 (top) and SUB (bottom). Dotted lines indicate bootstrapped 1st/99th percentile thresholds. **H)** Representational scale distributions for outbound and return routes. * denotes Kolmogorov-Smirnov test with P < 0.05. **I)** Scatterplot of CA1 (x axis) versus SUB (y axis) representational scale at identical locations for outbound (dark grey) and return (grey) routes. **J)** Mean representational alignment for the outbound (left) and return (right) runs for both CA1 (top) and SUB (bottom). Dotted lines indicate bootstrapped 1st/99th percentile thresholds. **K)** Variance in representational alignment for CA1 and SUB compared to bootstrapped 99th percentile. **L)** Mean derivative of the representational alignment for the outbound (left) and return (right) runs for both CA1 (top) and SUB (bottom). Dotted lines indicate bootstrapped 1st/99th percentile thresholds. Peaks (transition points) are locations of maximum change from reverse to forward representational alignment. **M)** Distributions of transition point distances relative to nearest turn apexes for CA1 (top) and SUB (bottom).

Both SUB and CA1 populations reliably encode individual locations at similar levels. Values along the diagonal of the correlation matrix correspond to even versus odd trial autocorrelations for the same location. If activity is consistent across trials at each location, correlations should be high (Cowen et al., 2014). This was the case for both SUB and CA1, as the median diagonal R values for both exceeded 0.7 (SUB = 0.76, CA1 = 0.74). The distribution of values for SUB was actually higher than that of CA1 (Figure 4C, one sided KS test, P < 0.0001, KS test stat = 0.17). Thus, both SUB and CA1 population activity patterns are reliable across trials at any single location, consistent with an encoding of location.

#### Representational scale

While both SUB and CA1 reliably encoded individual locations, the patterns in the correlation matrices suggest qualitative differences in spatial encoding. One feature of interest is the scale with which the two populations encode spaces. Consistent with historical precedent (Maurer et al., 2005), we operationalized the *representational scale* as the space surrounding a given location that is associated with population activity patterns that correlate at 0.5 or better to that location (Figure 4DE). This is determined by iteratively moving outward in both directions from any given location and finding the first position point at which the correlation value drops below 0.5. Notably, because task performance is associated with single directions of travel for all locations, drop-off points and their distance from any given location can be determined for the spaces visited both before and after the location of interest. The sum of these distances is the representational scale. Importantly, representational scale patterns were consistent for a wide array of correlation cutoff values (Supplemental Figure 2).

Representational scale of SUB was larger than CA1 (Figure 4F, one sided KS test, P > 0.0001, KS test stat = 0.37) with a median scale of 27 cm for SUB and 20 cm for CA1. This scale value was not constant across locations, however (Figure 4G). Both SUB and CA1 had larger scales on the return runs (Figure 4H, SUB: one sided KS test, P < 0.0001, KS test stat = 0.83; SUB: one sided KS test, P < 0.0001, KS test stat = 0.47). As a control to assess the validity of this dynamic change in scale, we created a bootstrapped distribution by randomly shifting the mean odd-trial and even-trial positional mean firing rate vectors together for each neuron independently. This procedure preserves field integrity and spatial information on a single neuron level but randomizes relationships between field locations and the environment. SUB representational scale on the return paths was often larger than predicted by the shuffled population, showing that the representational scale is dynamic in response to the environment features, even within one environment (Figure 4G, 117/197 positions of return outside 99th percentile of shifted population bootstrap). CA1 is on the edge of expected representational scale for return runs, showing a similar but muted scale dynamic (54/197 positions of return outside 99th percentile of shifted population bootstrap). The two regions scale together (Figure 4I, correlation of scatter, r = 0.49), suggesting a common underlying mechanism.

#### Representational alignment and segmentation to maze structure and task phase

Another stark feature of the correlation matrices was the presence of square regions of high correlations around the diagonals for both the CA1 and SUB correlation matrices (Figure 4A,DE). These regions are areas where the coding of the path is consistent, whereas their corners on the diagonal suggest locations that are spatially close but representationally more distant. To put it another way, the square regions indicate track regions that are chunked together and represented highly similarly.

To examine these regions of high correlation in quantitative detail, we found the difference in distances at which correlations drop off forward (to the right on the correlation matrix main diagonal) versus backward (to the left on the correlation matrix) at each location on the matrix diagonal and termed it the *representational alignment* (Figure 4DE). Positive values indicate forward-facing representational alignment relative to a specific location along a route in that a longer length of space ahead of the animal will exhibit similar population representation than the length of space behind the animal. Negative values indicate a backward-facing representational alignment in that a longer length of space leading up to a given track location will exhibit similar patterning than the spaces traversed subsequent to that location. A uniform alignment value would indicate a constant balance and a random distribution of place fields, whereas, alignments oscillating between positive and negative values indicate segmenting of space. Both SUB and CA1 dynamically changed their representational alignments as the animals navigated the maze (Figure 4J, CA1: 26% (89/337) of positions outside 99th percentile of shifted population bootstrap; SUB: 54% (182/337) of positions outside 99th percentile of shifted population bootstrap). Further, the variance of representational alignment for both regions was far greater than the shuffled population bootstrap (Figure 4K), suggesting there may be representational structure causing the large variations.

We therefore hypothesized that the dynamics of representational alignment followed the maze and route structure. Transition points at which the sign of the representational alignment flip from negative to positive indicate the end of one similarly represented section and the beginning of another. These transition points are the peaks of the derivative of alignment (Figure 4L). To ensure these variations are beyond expected random fluctuations if the place field distributions were random, only peaks that surpassed the 99th percentile of the shuffled population bootstrap were considered. Here, again, a major dissociation in between SUB and CA1 populations was observed. Transition points for the SUB population clustered at turns (Figure 4M, SUB circular median = 10.8°, P = 0.3877, circular median test, H_0_ = 0°). As turn locations can define the structure of the path network itself, this indicates that in SUB, population rate vectors exhibit persistent patterning over individual straight path segments. To our surprise, CA1 also transitions in forward-facing versus backward-facing representational alignment in CA1 population activity patterns. Unlike SUB, CA1 transition points were not near the apexes of turns, but instead surrounded the turns (CA1 circular median = 126.0°, P = 0.0023, circular median test, H_0_ = 0°). This distinction between SUB and CA1 shows a large-scale population difference in SUB and CA1 organization, and indicates that spatial representation in SUB cannot be considered as merely quantitatively different from CA1. Instead SUB and CA1 population activity patterns are organized differently relative to maze structure and task phase.

## Discussion

We compared spatial and directional tuning of CA1 and SUB neuronal populations during performance of a spatial working memory task within a complex, multipath environment. While a multitude of studies have documented the role of CA1 in location mapping and navigation, it has not been determined whether SUB functions as secondary output of the hippocampal formation with slight differences in the tuning properties or if SUB tuning to location and orientation can be considered a qualitatively different form of representation. That is, can SUB be regarded as appendage of the hippocampus or a unique node among a distributed set of brain structures encoding different types of organism-environment relationships. Here, using a multipath network that allowed us to compare many spatial and movement variables, we discovered that in three main ways, SUB exhibits striking differences from CA1. First, the spatially isolated firing fields for individual SUB neurons often occupied analogous locations across the environment, whereas CA1 neurons did not bear fields at locations with structural similarity. Second, while both SUB and CA1 population activity patterns are dynamic with respect to scale of representation, SUB scaling is considerably greater. Finally, SUB and CA1 populations differentially shifted in how they encoded contiguous locations behind versus in front of the animal’s current location. The recursion of SUB activity patterns at structurally analogous locations and the alignment of SUB population activity pattern shifts to the beginnings and endings of path segments evidences a role for this structure in encoding structurally-defined ‘chunks’ of the environment. Together, this evidence suggests that SUB may carry out a qualitatively different function during spatial navigation.

### Structural Analogy and Trajectory-Defined Discrimination Versus Generalization

Perhaps the most striking result from this study is evidence for representation of analogous, but spatially separate locations through recurrence of SUB firing patterns. By ‘analogous’ we refer to two or more locations that are spatially separate from each other, yet logically linked by reference to location within topologically similar pathways and/or by head orientations taken during travel through their locations. For humans, city street grids bear regularity in the orientations and intersections among individual pathways and, therefore, subsets of environmental locations have ‘analogy’ with respect to how they align relative to environmental boundaries. Thus, streets and avenues in major cities may be organized by north-south and east-west affordances for travel and give rise to emergent concepts such as a ‘northwest corner’ or ‘a block’. The observed spatial and directional firing patterns of SUB neurons and their organized recurrence across the maze used in our task is suggestive of the emergence of encoding that reflects recurrence in environmental sub-structure. Recurrence based on repetition within the structure of a single path has been reported for entorhinal cortex, retrosplenial cortex and posterior parietal cortex (Alexander & Nitz, 2015; Nitz, 2012; Derdikman et al., 2009) and the reported presence of ‘axis-tuned’ neurons in an earlier publication from our group (Olson et al., 2017) can be interpreted as reflecting recurrence of individual neuron firing patterns over all paths having the same orientation. We note that CA1 neurons under some circumstances exhibit tuning to related locations, but that, in the present case, CA1 neuron population patterns exhibited very little recurrence in firing patterns. Thus, the transition from CA1 to SUB would appear to reflect a substantive shift from encoding of location in the environment to the encoding of multiple locations according to structural and/or functional analogy.

Related to the phenomena described here are findings from earlier research showing that SUB generalizes in its spatial firing patterns across different environments sharing the same-shaped boundaries and/or a singular, prominent distal polarizing cue (Sharp & Green, 1994; Sharp, 1997; Brotons-Mas et al., 2010). In the present work, SUB neurons generalize in their spatial firing patterns across widely separated environmental locations that are analogous with respect to the shape and layout of paths taken through them. The analogous representations of SUB neurons sharply contrast with CA1 neurons. CA1 place cells are highly specific to experiences, segmenting experience by place, movement, and even trajectory dependence during navigation (O’Keefe & Dostrovsky, 1971; McNaughton, 1983; Markus & McNaughton, 1995; Wood et al., 2000; Frank et al., 2000; Ferbinteanu & Shapiro, 2003). Here, CA1 place cells, as previously reported, are highly specific to the animal’s location and trajectory. CA1 firing patterns clearly distinguished not only all separate locations within the environment, but also the first section shared by the four outbound paths according to whether the animal would ultimately turn left (paths 1,2) or right (paths 3,4).

The factors that lead to the SUB representations of analogous locations is an open question, but orientation appears to be an important variable. On our maze, analogous locations sharing highly similar SUB firing patterns are often aligned in orientation, although they may also have opposing directions such as the second segment of paths 1,2 versus paths 3,4 and the end segments of the two perimeter paths. The axis-tuned neurons previously reported from a minor sub-population of the present dataset (Olson et al., 2017) may be an extreme form of this sort of analogous representation. This may reflect association of inputs from different sub-populations of anterior thalamic neurons having opposite tuning by head orientation ((Taube, 1995; Goodridge & Taube, 1997; Peyrache et al., 2017; Viejo & Peyrache, 2019; Frost et al., 2020). Future work on environments where functionally similar spaces are not parallel may help tease apart whether the structural or functional aspects of the locations drive such responses in SUB neurons.

Previous work by Frank and colleagues (Frank et al., 2000) showed a high prevalence of path equivalency in deep entorhinal cortex neurons. These neurons fired in spaces that were analogous in their movement sequences, such as on left turns on similar sub-paths within a ‘W-shaped’ maze. The authors did not often see this pattern in CA1 neurons, however. Knowing both SUB and CA1 project to deep EC, the authors hypothesized SUB may play a role in this signal. Our data are consistent with this hypothesis and strengthen the possibility that analogous location representations are generated in SUB as a result of circuit interactions between EC, CA1, and anterior thalamic inputs that are tuned to head orientation.

An interesting quirk to our data exists in the trajectory dependence on the outbound routes. On the first segment leading into turn 1, CA1 population patterns almost completely differentiate trials in which the animal makes opposite choices at the upcoming turn. Even the SUB population demonstrates a much more moderate degree of trajectory dependence over the same locations. However, the segments leading to the final outbound turn (segment 3) show no trajectory dependence in either CA1 or SUB firing patterns. This difference could be due to the fact that the outbound path spaces subsequent to turn 1 are the same leading into both of the third turns (for paths 1,2 and 3,4). In contrast, different maze spaces lead into the first segment of all four outbound paths. We note here that the outbound turn 1 alternation pattern typically yields related alternation in the return paths leading into the first outbound segment.

Prior studies have either found strong evidence for trajectory-dependent modulation of place-specific activity (Frank et al., 2000; Wood et al., 2000; Ferbinteanu & Shapiro, 2003; Nitz, 2006) or the near absence of it (Bower et al., 2005). For multichoice mazes, prior work suggests that trajectory dependence was similar at the first and second choice points of a double-Y maze (Ainge et al., 2007; Grieves et al., 2016). Our data evidence the fact that trajectory dependence is dynamic in its expression even within a single environment and task structure. We hypothesize that the difference in its prevalence preceding different choice locations may be related to action stereotypy or to the navigational strategy employed by the animal. In our study, animals quickly traversed segment 1 and alternated at the first turn at a very high rate, 91%. However, the animals often slightly slowed preceding turn 3 and executed less regular turn choice patterns across trials. That is, the alternation patterns at turn 1 were not strongly observed at turn 3 such that different orders of paths to goal locations 1-4 were observed (Figure 2DE). We therefore suggest that the trajectory dependence is strongly related to relative degree of separation of the behavioral patterns. In other words, the trajectory dependence appears to follow the navigational strategy and patterns of the animals’ path selections in solving the task. Our data and associated hypotheses further predict that trajectory dependence should develop over learning and be more pronounced on tasks with more consistent transitions. We would predict that in our task, if path order was consistent, then trajectory dependence would appear for the locations leading into turn 3. If our theory is correct, it is also a potential indicator of the state of learning of the animal on a task.

### Representational Scale, Alignment, & Partitioning of Maze Segments

SUB neurons are known to have larger firing field sizes than those of CA1 neurons in the open field and on a track (Sharp & Green, 1994; Kim et al., 2012). It has been theorized that this may be useful for information transfer, as individual neurons carry more information (Kim et al., 2012; Kintashi et al., 2020). Using a population correlation approach (Maurer et al., 2005) we have shown that the SUB population also encodes space at a larger scale than CA1 - a result that follows from the larger fields of individual neurons.

While larger scale was an expected finding, we also report that the scale of CA1 and SUB is dynamic, changing on different segments of the track. For larger track segments, the encoding expands such that similar patterns are observed across locations separated by larger distances. SUB carries a larger dynamic range than CA1 in this respect, but there appears to be parallel shifts in scale of representation between the two regions. This suggests a shared underlying cause. Differences in CA1 and CA3 scale are only known to exist across the longitudinal axis (septo-temporal axis) of the hippocampus (Maurer et al., 2005; Jung et al., 1994; Kjelstrup et al., 2008; Royer et al., 2010). This has been theorized to be due to a differential relationship between place fields and running speed across the longitudinal axis (Maurer et al., 2005). We therefore hypothesize that the increased running speed on longer track sections may be responsible for the dynamic scaling reported here, but emphasize that this influence can apply to the same population of neurons as opposed to expression only across the longitudinal axis.

We also examined the locations at which spatial firing patterns in SUB and CA1 shift. From these analyses, we find that CA1 and SUB population firing patterns partition the environment in different ways with SUB population shifts biased to the locations of path intersections. The partitioning we observe is reminiscent of “chunking”, a proposed psychological phenomenon where information is grouped into intuitive components to facilitate memory (Miller, 1956). In our data, there are transition points of low representational scale separating areas of large scale. Before and after these transition points, SUB and CA1 patterns may persist across contiguous locations along the track. We note that, in such cases, the patterns preceding and following a transition point are different, and that this, in part, defines the transition point itself. Previous research has suggested SUB neurons’ activity results in a schematized chunking of space (Olson et al., 2017; Lee et al., 2018), and our results largely support this interpretation. Gupta et al. (2012) previously described similar dynamics in chunking for CA1 populations relative to inflection points in the animals behavior (e.g., turns and stopping points for reward). Our results complement their findings, showing that the activity patterns segmenting space in this fashion are prevalent even at a population level, occur with less frequency for SUB than for CA1, and, as stated, are biased to path intersections for SUB.

If chunking is indeed the outcome of spatially organized, non-continuous shifts in spatial representation, then an important question is how the information (space) is actually being grouped. This may lend insight into processing of the task and space. Instead of presupposing the locations, we applied an unbiased analysis of all potential locations for transition between representational partitions (or “chunks”). Perhaps surprisingly, we found the transition locations were not the same for CA1 and SUB. SUB transition points near shifts between path segments (i.e., at corners) were consistent with the importance of orientation previously discussed. CA1 transitions, on the other hand, often occurred within individual path segments, for example, the halfway point along segment 1 of outbound paths. These results may hint at the different functions of the two hippocampal outputs. The SUB, with its more orientation-based partitioning of maze space, may be more specific to structural components of the full path network while for CA1 transition may more closely align to action sequences.

To report that SUB is both chunking space and representing analogous spaces may seem superficially contradictory, but in many ways, the two features are orthogonal. Analogous representations treat spatially separate locations similarly, while chunking groups locally contiguous space. Together, both encoding features function to assemble and associate particular kinds of places together. Indeed, progress along a learned route of a particular shape, irrespective of its location in an environment, is encoded independently of action in many posterior parietal and retrosplenial cortex neurons (Nitz, 2006; Alexander and Nitz, 2015). SUB activity patterns may thus contribute to such encoding through input patterns to retrosplenial cortex that recur for topologically similar, but spatially separated routes.

### Encoding of Location

As with any set of neural analyses, the importance of the relative aspects of CA1 and SUB activity patterns depends on the transformation of information by downstream readers. Here, we have analyzed population activity using correlations of 0.5 as a baseline due to historical precedence (Maurer et al., 2005). While our results on representational scale, alignment, and chunking are robust and similar at a variety of thresholds between 0.3 and 0.7 (Supplemental Figure 2), what threshold is meaningful is ultimately determined by the neural networks receiving the signal.

The implications of what level of correlation is meaningful is especially important to SUB outputs, as differing levels will determine if SUB generalizes or separates locations sharing spatial variables. Odd/even trial population correlations at individual locations were extremely high in SUB and, if anything, higher than for CA1. The differences between these regions lies a step below the strong “place” correlations in population patterns for repeated visits to the same location while on the same trajectory. Thus, if downstream readers are extremely sensitive to small gradations in firing rates among a large population of neurons, many locations will therefore be differentiated and an encoding of location only may result. However, if output regions are less sensitive to variations in input patterns, the analogous locations will be read equivalently, and structural information will be the information transmitted. We cannot determine, as yet, whether structures such as retrosplenial cortex better resemble the location tuning of CA1 as opposed to the location and analogy encoding properties of SUB. However, we predict that population coding for different, but analogous locations in SUB often reaches values of 0.75 or higher, levels often considered adequate evidence for reliable encoding of individual locations (e.g., Maurer et al., 2005).

### Anatomical Context

Considering the importance of downstream readers, it is worth considering outputs of SUB and their known activity patterns in light of these novel input signals from SUB. SUB projects to many regions considered important for spatial navigation. One particular region of interest may be retrosplenial cortex. Retrosplenial cortex primarily receives its “hippocampal” input from SUB (Jay & Witter, 1991) and, like SUB, has been shown to also represent conjunctions of spatial and movement variables (Alexander & Nitz, 2015). The conjunctions seen in RSC often combine allocentric with egocentric variables such as proximity to a border and locations of turns (e.g. Alexander et al., 2019; Alexander and Nitz, 2015), an aspect not seen in SUB to date. This may further indicate that SUB acts in part as a step before retrosplenial cortex along the transformation of spatial information into action.

Finally, these results throw into doubt the idea that the activity patterns in SUB are best understood as a mere transformation of CA1. If this is not the case, it raises the question as to what the other key drivers of the activity in SUB are. Strong cortical inputs exist from entorhinal, perirhinal, and postrhinal cortices and subcortical inputs from nucleus reuniens, the medial septum, and anterior thalamus (Witter et al., 1990; Witter et al., 2000; O’Mara, 2006; Witter, 2006). Recent work has shown the importance of anterior thalamic inputs into SUB in guiding choice behavior at intersections. Temporary and permanent anterior thalamic lesions led to marked behavioral deficits and degraded spatial firing in SUB while CA1 place fields remain intact (Frost et al., 2020). Further research is needed into investigating the importance of these inputs for SUB function.

### Limitations and Future Directions

While our task and data have brought to light many new interesting aspects of CA1 and SUB activity during spatial navigation, there were aspects of the experiment we would have liked to improve. We believe the Triple T maze is a strength of our design, allowing assessment of multiple spatial variables, their conjunctions, and the ability to assess generalization. That said, in order to more conclusively tease apart contributions of variables that are highly conjunctive, even more data and combinations are needed. Including all 8 reward locations and repeating the task in both directions would have moved us much closer to this goal, but this amount of data is largely infeasible in single recording sessions. Future work may need to record animals over multiple sessions in a day to collect the varied behaviors needed to better disentangle the forms of information represented in SUB.

Another limitation of our work is the anatomical distribution of our recordings. Most of our data was collected in proximal SUB. The proximal-distal gradient in SUB is well described and separates both activity patterns, sources of afferents, and projection targets (Naber & Witte, 1998; Kim et al., 2012; Aggleton & Christiansen, 2015; Cembrowski et al., 2018). It is of note that this data, while appearing extremely spatial, is in the region of SUB that largely projects to and from lateral entorhinal cortex, not medial. Conversely, the object vector cells recently discovered in SUB appear to have been located predominantly in distal SUB (Poulter et al., 2020). This leads us to speculate that the what/where division currently dominant in the field between lateral and medial entorhinal cortex may be a false dichotomy, and that in the context of more complex behaviors, lateral entorhinal cortex activity may be interpreted differently. Regardless, future work spanning the full proximal-distal extent of SUB on this or similar navigation tasks would be invaluable toward having a better understanding of the proximal-distal function of SUB for spatial navigation.

## Conclusion

This work has studied the properties of neurons in dorsal CA1 and SUB while rats navigated a complex path network. We have found that both CA1 and SUB dynamically adapt their encoding properties over different phases of the task but chunk those spaces in different locations. SUB neurons showed a propensity for activity at analogous functional or structural locations on the track, and the SUB population often encoded these analogous locations similarly. The differences in specificity and generalization between these two hippocampal outputs are stark and suggest SUB and CA1 may play complementary roles. Differentiating small differences as seen in CA1 is crucial for memory of individual events, but grouping similar information as in SUB is vital for extrapolation and a holistic cognitive map. As such, we believe these results point to complementary roles for CA1 and SUB in episodic memory and navigation, and that these roles may expand more generally to episodic storage versus consolidation of abstract information. We hope future work will further evaluate these two hippocampal outputs side by side to clarify their roles in navigation and memory.

## Methods

### Subjects

Subjects were adult male Sprague-Dawley rats (N = 6). Rats were housed individually on a 12-h light/dark cycle. Prior to experimentation, animals were habituated to the colony room and handled for acclimation for 1-2 weeks. The rats were food restricted for motivational purposes at 85-95% of their free-fed weight. No water restriction occurred. All rats were required to reach a weight of 350g (5-10 months of age) before surgery and experimentation. All experimental protocols adhered to AALAC guidelines and were approved by UCSD IACUC and UCSD Animal Care Program.

### Apparatus

All behavioral tasks were conducted on the “Triple-T” track maze. The Triple-T maze is a custom built environment made of black plastic with a running surface of a thin sheet of black rubber. This path-network is made of tracks that are 8cm in width. The overall dimensions of the environment are 1.6m x 1.25m and are elevated 20cm from the ground. Track edges are approximately 2cm in height, allowing the rat an unobstructed view to distal cues. Taller 10cm walls were included at internal sides of corners and to block potential shortcuts across small gaps between tracks near the reward locations.

### Behavior

Rats were habituated to the Triple T maze with two 30 minute epochs of free exploration with rewards scattered throughout the maze. Animals were then trained to conduct outbound routes, traversing from the midpoint of one of the long edges of the maze (Figure 1A, location of rat), through the center space, to one of the four reward locations (Figure 1A, colored dots). The outbound routes from the start to the reward locations consisted of straight paths interleaved with 3 left or right turns. The first and third turn directions were choice points but the second turn was forced. Total outbound route lengths were 140cm with turns at 51cm, 87cm, and 118cm. Rewards (1/4 -1/2 piece of Cheerios cereal) were manually delivered at the reward sites. Animals were trained to return to the start location via the outside paths after completion of an outbound route. We refer to these behaviors throughout as return routes. Animals were trained over 1-2 weeks to make these route traversals and to do so without stopping mid-route. At this point in training, we implemented the visit-all working memory task to be used throughout experimentation. The animals were rewarded at any of the four locations, but needed to visit all four locations before rewards were again available at previously visited reward locations. Over at least 2 additional weeks, animals were trained by simple trial and error to a criterion of 80% for ballistic (uninterrupted) path traversals, regardless of reward performance. Only after this level of running performance were animals surgically implanted for experimentation.

### Surgery

Rats were surgically implanted with 1-3 tetrode (twisted sets of four 12μm polyimide-insulated tungsten or nichrome wires) arrays integrated into custom microdrives. Each microdrive held four to twelve tetrodes. Under isoflurane anesthesia, animals were positioned in a stereotaxic device (Kopf Instruments). Following craniotomy and resection of dura mater, microdrives were implanted. Microdrives were implanted relative to bregma with targeting coordinates consistent across animals. SUB: A/P −5.6 to −6.6 mm, M/L ±1.6 to ±2.7 mm; dorsal CA1: A/P −3.8mm, M/L ±2.3mm.

### Neural and Behavioral Recordings

Following a week for recovery from surgery, animals were retrained for at least 1 week before beginning recordings. This ensured adequate behavior and running ability with the new weight of the implant. Electrodes were moved ventrally in 40μm increments between most recordings to maximize the number of distinct units collected for each animal. Each microdrive had 1-3 electrical interface boards (EIB-16, Neuralynx) connected to a single amplifying headstage (20x, Triangle Biosystems). The headstages were tethered to a set of preamplifiers (50x) and a high pass filter (>150 Hz). These signals were input in the acquisition computer running Plexon SortClient software. Signals were recorded after digital filtering at 0.45–9 kHz, 1–15x amplification (to reach a total of 1,000–15,000x amplification of the signal), and 40 kHz digitization. Waveforms from single units were isolated in Plexon OfflineSorter software. Waveform parameters used were peak height, peak valley, energy, average voltage, full width at half maximum, and principal components. Waveform clusters appearing to overlap with the amplitude threshold set for collection were discarded to avoid collection of only partial spiking data of neurons. Waveform amplitudes were monitored to ensure systematic fluctuation did not result in confounds for isolating single units. Recordings typically lasted 30-60 minutes.

After single unit isolation was complete, cluster quality was quantified using a modified isolation distance score as described in Olson et al. (2017, Figure 2B). Briefly, isolation distance (Harris et al., 2000) measures the cluster separation using the Mahalanobis distance between the cluster center and the *n*th closest noncluster spike, where *n* is the number of spikes in the cluster. The units of this are equal to cluster variance. Equivalently, isolation distance is the radius of the cluster center to the circle containing double the number of spikes as actually classified in the cluster. This is normalized by the cluster variance. As such, this measure is undefined when the spikes recorded is not at least double the spikes included in the cluster. To avoid this issue, Olson et al (2017) adapted this measure to be the minimum of the isolation distance as defined by Harris et al. and the distance to the noncluster spike 20% into the noncluster spike distance distribution. This modification defines the value for all neurons and reduces the isolation distance for clusters with many spikes. This modification makes the criteria stricter, but this conservative adjustment more accurately represents cluster quality for tetrodes with one high firing neuron. Isolation distance is not a criterion used for excluding units in this study. Instead, it is presented here to show the high quality of neurons identified.

Animals’ positions were tracked during neural recordings using a camera located 2.6m above the recording room floor. Plexon CinePlex Studio software detected two LED lights on the animal’s surgical implant separated by approximately 5cm. Location tracking was captured at 60Hz. At any given time point, position and orientation were determined using the average location of the two lights and the orientation of the vector between the lights. All animal movement data such as location, head direction, and derivatives are calculated from these values.

We recorded a total of 573 subiculum and 401 CA1 neurons from 6 rats. Relatively inactive neurons were excluded from analysis if their activity never averaged at least 3 spikes/sec at any location on the maze. Interneurons were attempted to be excluded from analysis by removing neurons that averaged more than 3 spikes/sec at all locations on the maze. After exclusions, our dataset consisted of 480 subiculum and 298 CA1 neurons.

### Histology

Animals were perfused with 4% paraformaldehyde under deep anesthesia. Brains were extracted and sliced into 30μm -50μm sections. Slices were Nissl stained to reveal the final depth of electrodes. Microdrive movement was used to reconstruct recording-depth profiles. These data were then compared to known anatomical boundaries of the regions of interest (Paxinos & Watson, 2006) to establish a final categorization of the source of each microdrive.

### Filtering of Behavior for Fluid Route Traversals

To limit variation in behavior at each location, we only analyzed data for when the animal was performing a smooth, uninterrupted traversal of an entire route. We refer to these smooth traversals as uninterrupted trials. This permitted us to examine action potential firing rate data associated with stereotypical movements (forward running, turning) through all sections of all routes. The defined routes were the four internal outbound routes toward potential reward sites and the two return routes leading back to the internal entrance (Figure 1D).

A multistep process using custom MATLAB graphical user interfaces was used to identify uninterrupted traversals of each route. Users defined starting and ending gates for each route. The program then selects all trials crossing these locations with sustained running speeds of 3cm/s or greater throughout. Finally, a researcher verifies all identified trials did not contain either obvious interruptions in locomotion or significant deviation from the stereotyped path for that route (Figure 1D). This method separates stalled track traversals, reward periods, and behavior between trials from controlled action and spatial data. Individual recordings were excluded from analysis if the number of traversals on each internal route did not exceed 3.

### 2D Positional Firing Rate Maps

To assess activity as a function of 2D space, we calculated individual neurons’ positional firing rates by dividing the total number of spikes at each location by the total occupancy time at that location. If only showing identified routes, only data identified as during an uninterrupted trial was used. Positional firing maps were smoothed using a 2D convolution with a Gaussian filter with s.d. of 1cm that also accounts for bins with no occupancy (Kraus, 2013).

### Spatial Statistics

Many spatial statistics have been previously defined to summarize the spatially selective nature of neuron firing. First, we created 2D positional firing rate maps using 1cm square bins with no smoothing. We then calculated spatial information in bits per spike (Skaggs et al., 1993),

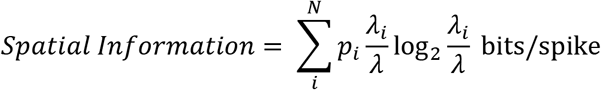

where *i* is each bin, *p*_*i*_ is the occupancy of bin *i, λ*_*i*_ is the mean firing rate of bin *i*, and *λ* is the overall mean firing rate. Spatial selectivity (Skaggs et al., 1996) is large for neurons with firing fields that are a small portion of the environment and is simply the ratio of the max firing bin divided by the overall mean firing. Finally, spatial coherence (Kubie & Muller, 1989) assesses continuity of signals and is defined as the z-transform of the spatial autocorrelation between each bin and the average of its immediate neighbors. Here we used a Spearman correlation as the correlation and do not transform into Z coordinates.

### Linearized Firing Rate Maps

We also aligned neural activity to progression through each of the identified routes. For each recording, custom MATLAB software is utilized to generate a spatial template matching the average trajectory of the animal along each route in the horizontal (2D) plane. This approach ensures the best possible matching of animal behavior and positions taken across recordings and trials. Position samples included in the identified ballistic route traversals were mapped to the nearest template bins. Then the linearized firing is found by dividing the total number of spikes at each location by the total occupancy time at that location. Linearized firing maps were smoothed using a 1D convolution with a Gaussian filter with s.d. of 1cm that also accounts for bins with no occupancy (Kraus, 2013). The linearized firing rate is then averaged over all traversals the animal made for that recording to calculate the mean linearized firing rate.

### Individual Neuron Analogous Route Correlations

To assess individual neuron’s similarity of representation across routes, a pairwise Pearson’s correlation was calculated for each neuron between each outbound route and each other outbound route. The calculation was on the vector of mean linearized firing rates between the non-overlapping portions of the routes. The same analysis was then applied to the two return routes. The maximum correlation of all correlations was used for the population analysis, as the sparse nature of activity in HPC and SUB renders most values undefined or near-zero. Of interest is whether any high correlations exist among any of the comparisons.

### Population Analogous Route Correlations and Correlation Matrices

Population-wide similarity of route representations were also taken. For each position along the linearized track space, individual neurons’ mean firing rates were appended to create a population firing rate vector. Then, Pearson’s correlations were calculated of the population firing rate vectors of two locations of interest. Locations chosen include locations with similar but spatially separated locations or functions with respect to the trained behavior (i.e. two return routes, or close to the reward locations). If more than two analogous locations existed, pairwise correlations were calculated and then averaged across all pairwise combinations.

As a control, the linearized firing rates for each neuron were split into odd and even traversals. For each neuron a new mean firing rate was calculated from the odd/even traversals and used to create two population firing rate vectors from non-overlapping trials. Correlations of the same space from these two odd and even vectors show the reliability of the signal and serve as a control for the limit of possible correlation given the consistency patterns of the population.

Correlation matrices were also constructed to visualize the relationship of each pairwise set of positions across the linearized track space. As specified above, the linearized firing rates for each neuron were split into odd and even traversals. For each neuron a new mean firing rate was calculated from the odd/even traversals. For each pairwise combination of positions, the population vector of odd traversal mean firing rates was correlated with the population vector of even traversal mean firing rates. By splitting the trials into non-overlapping sets and correlating these values, the diagonal value of the correlation matrix reflects a form of reliability of population activity for each position.

### Population Representational Scale, Alignment, and Transition Points

The correlation matrices exhibit where and how similar population neural activity patterns are across the maze. We developed new and adapted old metrics to quantify some of these patterns.

*Representational scale* was defined as the space surrounding a location that correlates above a given threshold with that location. This groups contiguous, similar space both in front of and behind the location. We used 0.5 as the correlation threshold, consistent with historical precedent (Maurer et al., 2005), for this analysis and its derivative analyses below, but a wide array of values resulted in similar results (Supplemental Figure 2). *Representational alignment* was operationalized as the difference in the amount of highly correlated space before and after the current location. More correlated space in front of the location is positive, while more space behind is negative. The derivative of the representational alignment shows how the alignment changes across the track space. The peaks of the derivative, termed the *transition points*, are points where the correlations move from backwards looking to forwards looking, indicating a separation in the representation of nearby locations.

### Representational Analyses Shifted Population Bootstrap

To assess the magnitude and validity of the fluctuations of our newly defined representational scale and alignment metrics, we created a bootstrapped distribution for comparison. This distribution was created by appending all of the routes linearized firing rates, as done in the correlation matrix, then shifting the mean firing rates of individual neurons independently to create a new population where local neural activity field characteristics remain, but population level neural field distributions are random and unassociated with track features. New correlation matrices were created from this shifted population and all representational analyses were conducted. This procedure was repeated 1000 times, and 1st and 99th percentiles of the results were used as statistical comparisons for results expected by similar neurons with similar field properties by chance.

### Statistical Tests

Nonparametric tests were used throughout to avoid assumptions of normality in the data. The Mann Whitney U test was used to evaluate the statistical significance of behavior to chance and differences between SUB and CA1 across the different analyses, as well as comparisons of SUB and CA1 to odd trial or even trial control populations. The Kolmogorov-Smirnov test was used to assess if pairwise distributions of correlations from SUB and CA1 were significantly different. Representational scale, alignment, and transition points were compared to the 99th percentiles of a population created by bootstrapping the shifted linearized firing rates from the actual neural population. Circular median tests were used to compare CA1 and SUB transition points to the turn locations. No statistical methods were used to predetermine sample sizes. However, based on similar sample sizes reported in previous publications, we believe we have adequate power (0.8) or greater to detect significant effects.

## Data and Code Availability

The data collected and analyzed in this study as well as the code used in all analyses are available from the corresponding author upon reasonable request.

---

**Supplemental Figure 1:**
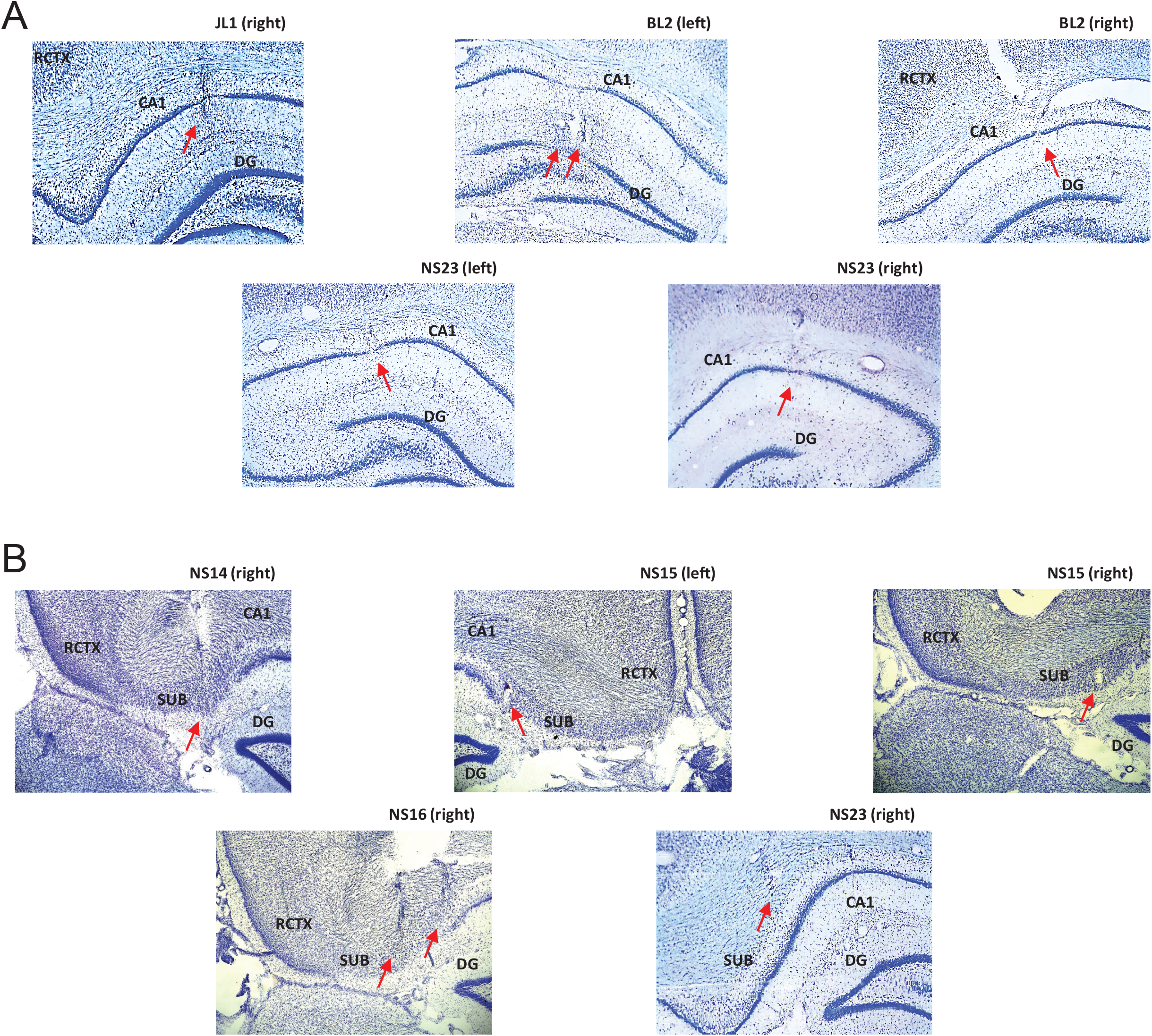
Summary of recording site histological data. **A)** Recordings of CA1 neurons (N = 401) were obtained from a total of six four-tetrode bundles in three animals. Numbers of total recorded neurons and numbers of neurons included above each figure. Red arrows depict tracks left by the bundles and their approximate endpoints. All of the recording sites were restricted to the dorsal CA1 while one (BL2-right) was additionally moved ventral to DG after main-experiment (DG data not included). **B)** Recordings of SUB neurons (N = 573) were obtained from a total of six four-tetrode bundles in four animals. Red arrows depict tracks left by the bundles and their approximate endpoints. Three of the recording sites were restricted to the SUB while three (NS15-left, the lateral bundle in NS16-right, and NS23-right) were in a transition zone bordering the CA1 sub-region. **Abbreviations: RCTX (retrosplenial cortex), DG (dentate gyrus), SUB (subiculum)**

**Supplemental Figure 2:**
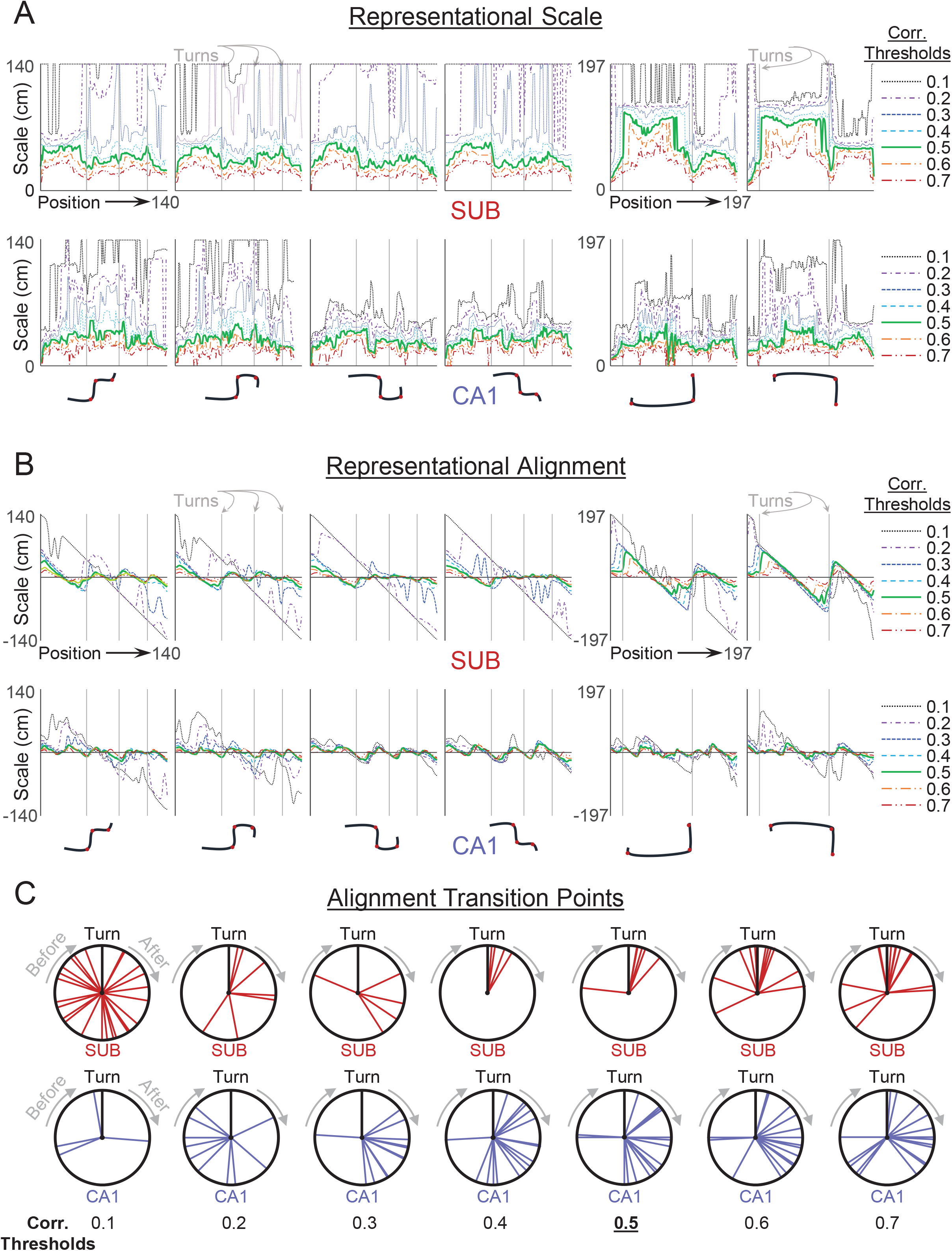
Representational Scale, Alignment and Transition Points are Consistent across a Wide Range of Correlation Threshold Values. **A)** Representational scale as a function of correlation thresholds for each of the four outbound and two return runs. *Top Row:* SUB population. *Bottom Row:* CA1 population. **B)** Representational alignment as a function of correlation thresholds for each of the four outbound and two return runs. *Top Row:* SUB population. *Bottom Row:* CA1 population. **C)** Representational alignment transition points as a function of correlation thresholds. *Top Row:* SUB population. *Bottom Row:* CA1 population.

